# Harmonized nucleoside mass spectrometry enables reproducible cross-platform RNA modification quantification

**DOI:** 10.64898/2026.07.17.739095

**Authors:** Jan Felix Dalwigk, Kira Kerkhoff, Oskar Knittelfelder, Robert L Ross, Maria Cristina Petrella, Anna Kuśnierczyk, Tulsi Bhandari, Aurore Attina, Alicia Burkard, Carolina Brás-Costa, Michael S. DeMott, Kitty Johnson, Xenia Kerkhoff-Bernaciak, Jennifer Kist, Ken Kögel, Martina Krämer, Ricardo Moreno-Ballesteros, Ganna Podoprygorina, Kaley Simcox, Özge Simsir, Anton Skriba, Sam Wein, Hana Cahova, Benjamin A. Garcia, Mark Helm, Alexandre David, Katharina Höfer, Sebastian Leidel, Patrick A Limbach, Eva Novoa, Eduard Sabidó, Sabine Schneider, Philippe Wolff, Peter Dedon, Vivian Cheung, Stefanie Kaiser

## Abstract

RNA modification analysis by LC–MS/MS is central to epitranscriptomics, yet quantitative comparison across laboratories and instrument platforms remains poorly standardized. Here, we performed a community-driven benchmarking study during the first Human RNome Project workshop to systematically evaluate cross-platform reproducibility of ribonucleoside mass spectrometry workflows. Using the same analytical column and gradient, standardized RNA samples, and shared reagents, we compared nucleoside quantification across quadrupole, time-of-flight, and orbitrap-based LC–MS platforms employing distinct acquisition strategies.

While chromatographic separation was highly reproducible across systems, nucleoside-specific MS response behavior differed substantially between platforms and limited direct comparability of relative signal intensities. These response differences varied across analytes and concentration ranges, demonstrating that harmonized chromatography alone is insufficient for transferable quantitative analysis. Stable isotope-labeled internal standard (SILIS) normalization substantially reduced platform-and method-dependent response and improved agreement for most evaluated modifications. External calibration improved agreement between qTOF and Orbitrap workflows for a subset of modifications but did not fully resolve residual intersystem differences.

Based on these findings, we establish benchmark-derived recommendations for harmonized relative and absolute RNA modification quantification, including guidance for calibration design, quality control, and data reporting. Together, this work provides a methodological framework for reproducible nucleoside LC–MS/MS workflows and establishes a foundation for large-scale comparative epitranscriptomic studies.

**Graphical abstract:** 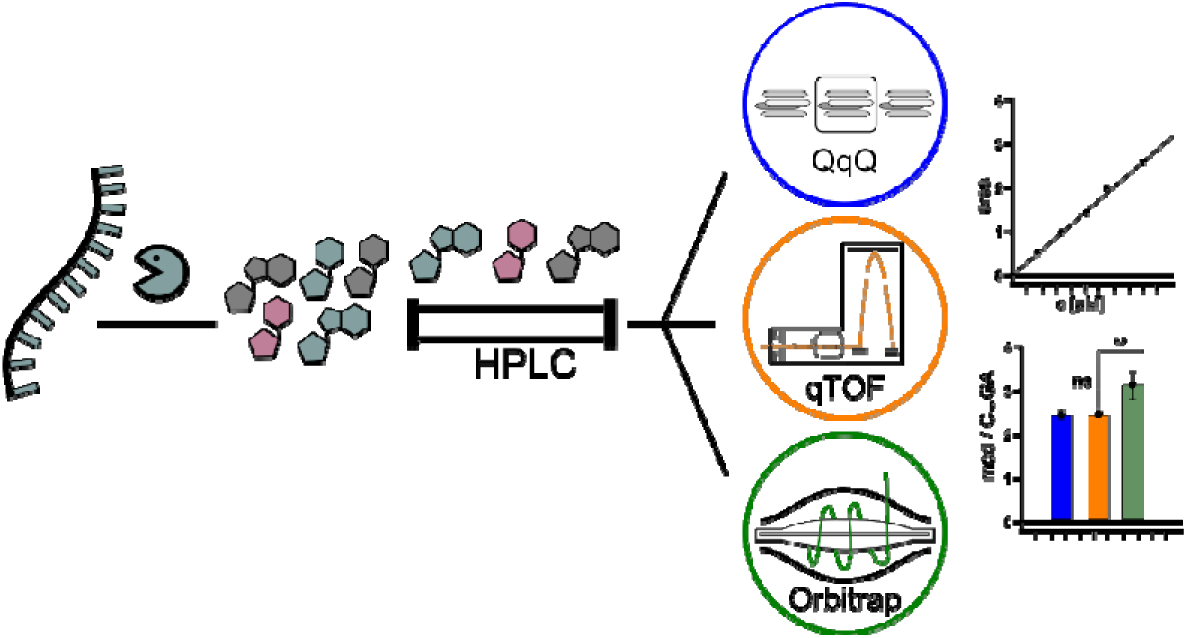

## Introduction

To date, more than 170 naturally occurring RNA modifications have been identified (1–5). Many of these modifications are recognized to affect RNA stability, structure, decoding fidelity, and RNA–protein interactions, which makes them central regulators of gene expression. Dysregulation of RNA-modifications is linked to the pathogenesis of at least 100 human diseases (6–8). Therefore, characterization of these modifications has become a central objective in epitranscriptomics. Due to its sensitivity, molecular specificity, and broad modification coverage, liquid chromatography coupled to mass spectrometry (LC-MS) remains the gold standard for direct chemical identification and quantitative analysis of modified ribonucleosides. For quantification, nucleoside MS is the most common approach, because it allows for sensitive and accurate analysis of complex RNA pools. Here, the RNA is enzymatically hydrolyzed into single nucleosides. This converts the potentially broad and complex variety of different RNA sequences into a nucleoside pool with fewer distinct species, thereby facilitating straightforward quantification by LC-MS/MS. Absolute and relative quantification approaches must be further distinguished within nucleoside MS, as they address fundamentally different biological and analytical questions and pose different challenges for validation and harmonization (Table 1).

**Table 1.** Quantification types in nucleoside mass spectrometry of the epitranscriptome.

| Quantification type | Biological Question | Primary analytical requirement |
| --- | --- | --- |
| Relative quantification | Modification changes between different conditions | Stable intra-system conditions<br>Signal linearity |
| Absolute quantification | Modification stoichiometry | External calibration<br>Consistent concentration:signal relationship |

For relative quantification the instrument’s MS response of a modified nucleoside is evaluated. To control for differing input amounts, this MS response must be normalized. Normalization can be achieved by relating the MS response of the modified nucleoside to either the UV area (9) or the MS response of canonical nucleosides (10,11). Finally, this resulting ratio can be compared between different samples within a sample set and reported as a fold change. Relative quantification can be used to compare modification frequencies between RNA classes and assess changes in RNA modification composition during cellular differentiation, stress adaptation, or disease progression. Furthermore, it is used to identify dynamic modification changes associated with homeostatic perturbations, drug treatment, or genetic manipulation. The numeric value of normalized response is arbitrary and largely platform dependent, so biological interpretation of acquired results can be limited.

Stoichiometric values can be obtained through absolute quantification. Here, the abundance of a modified nucleoside is determined in defined physical units, such as fmol per µg RNA or modifications per cell. Absolute quantification is primarily used to estimate modification stoichiometry, compare modification levels across studies or laboratories, and monitor global changes in RNA modification abundance under defined physiological or pathological conditions. While providing a valuable additional layer of information, the use of nucleoside standards in known concentrations is required. Thus, absolute quantification is often limited by the high costs and restricted availability of synthetic nucleoside standards. In addition, absolute quantification requires analyte-specific calibration models that are experimentally validated with respect to their applicable concentration range, regression behavior, and quantitative performance. These requirements fundamentally distinguish absolute from relative quantification and represent a major challenge for harmonized cross-platform analyses. In summary, these two distinct analytical objectives differ substantially in their requirements for calibration and quantitative transferability across analytical workflows and mass spectrometry platforms.

Recent harmonization efforts within the Human RNome Project demonstrated that robust inter-laboratory reproducibility can be achieved for targeted triple quadrupole-based nucleoside workflows despite differences in sample preparation, RNA hydrolysis, and instrumentation (12). Different laboratories currently employ various MS response correction strategies for nucleoside MS such as the use of single internal standards (9,11), stable isotope-labeled internal standard (SILIS) mixtures (13,14), or no internal standard (15). In parallel, substantial variability exists across the broader analytical workflow, including RNA processing, enzymatic digestion, chromatographic separation, instrument configuration, acquisition strategy, and downstream data analysis (10,16). Previous harmonization efforts and methodological studies have highlighted many of these sources of variability and identified critical pitfalls associated with RNA modification analysis, including chemical instability of specific ribonucleosides such as N4-acetylcytidine (ac^4^C) (17) and enzymatic or chemical artifacts (18–21) including adenosine deamination during sample handling (10). Despite these advances, the extent to which such methodological differences influence quantitative transferability across fundamentally different mass spectrometry platforms remains poorly understood.

Previous benchmarking efforts primarily focused on targeted triple quadrupole workflows (12). However, the rapidly expanding epitranscriptomics field increasingly relies on a broader range of analytical architectures, including orbitrap- and time-of-flight (TOF)-based systems employing distinct acquisition strategies, detector designs, and signal processing approaches (22–24). These platform-specific differences may introduce analyte-dependent quantitative biases by affecting ion transmission, fragmentation behavior, detector response, and dynamic range. Thus, reproducible quantification across laboratories does not necessarily translate to consistent quantification across fundamentally different mass spectrometry platforms. At present, it remains unclear to what extent nucleoside quantification can be directly transferred across analytical platforms or which analytical strategies are required to achieve robust cross-platform agreement. This lack of harmonization currently limits comparability between datasets generated across laboratories and complicates the establishment of reproducible large-scale epitranscriptomic studies.

Here, we performed a harmonized cross-platform benchmarking study during the first Human RNome Project workshop to systematically evaluate the transferability of nucleoside MS analysis across distinct mass spectrometry architectures. Using standardized RNA samples, shared reagents, harmonized chromatographic conditions, and common calibration strategies, we compared quantitative performance across triple quadrupole, orbitrap, and time-of-flight platforms employing different acquisition principles. We assessed chromatographic reproducibility, platform-dependent response behavior, and compared relative and absolute quantification strategies. Our analyses demonstrate instrument-specific response behaviors that can be partially mitigated through external calibration and largely resolved by employing stable isotope-labeled internal standards. Finally, we integrate the workshop results and recent recommendations for nucleoside analysis into an end-to-end workflow guideline.

## Materials and Methods

### List of chemicals and reagents

Unless stated otherwise, all chemicals and reagents were purchased from Sigma-Aldrich (St. Louis, MO, USA).

For the Agilent (Santa Clara, USA) and Bruker (Billerica, USA) instruments, ultra-LC-MS grade acetonitrile was purchased from Carl Roth (Karlsruhe, Germany) and the aqueous phase was prepared using ultrapure water produced by a Milli-Q Integral system (Merck Millipore, Darmstadt, Germany). For the Thermo Fisher Scientific (Waltham, USA) system, both acetonitrile and water were purchased from Thermo Fisher Scientific (Waltham, USA) and provided in clear glass bottles. The nucleoside standards Pseudouridine (Ψ), 3-(3-amino-3-carboxypropyl)uridine (acp^3^U), 1-methyladenosine (m^1^A), 7-methylguanosine (m^7^G), 5-methylcytidine (m^5^C), 2‣-*O*-methylcytidine (Cm), 5-methyluridine (m^5^U), 2‣-*O*-methyluridine (Um), 2‣-*O*-methylguanosine (Gm), 2‣-*O*-methyladenosine (Am), 6-methyladenosine (m^6^A) were obtained from Biosynth (Staad, Switzerland). 3-methylcytidine (m^3^C), 1-methylguanosine (m^1^G), 5,2‣-*O*-dimethyluridine (m^5^Um), *N*^6^-threonylcarbamoyladenosine (t^6^A), *N*^2^, *N*^2^-dimethylguanosine (m^2,2^G) were purchased from TRC (Toronto, Canada). 3-methyluridine (m^3^U) and 2-methylguanosine (m^2^G) were obtained from Benchchem (Ontario, USA). Inosine (I) was obtained from Carbosynth (Newbury, UK), *N*^6^-isopentenyladenosine (i^6^A) from ChemScene (Monmouth Junction, USA), *N*^6^, *N*^6^-dimethyladenosine (m^6,6^A) from Alfachemistry (New York, USA), Dihydrouridine (D) from Apollo Scientific (Stockport, UK) and *N*^6,2^‣-*O*-dimethyladenosine (m^6^Am) from Berry and Associates (Atlanta, USA).

#### Cell culture and total RNA extraction

Human B-cells were obtained from Coriell Cell Repositories (GM12878). Cells were cultured in RPMI 1640 medium (with L-glutamine) supplemented with 15% heat-inactivated fetal bovine serum (FBS) and 1% Penicillin/Streptomycin at 37°C and 5% CO_2_, and harvested at 1×10^6^ cells/ml. RNA was extracted using Qiagen RNeasy columns (Qiagen, catalog #75144) following the manufacturer’s protocol and eluted from the column in two washes of 250 µL diethylpyrocarbonate (DEPC)-treated water (Thermo Fisher Scientific #AM9915G; autoclaved by the manufacturer to remove residual DEPC). RNA concentration and integrity were assessed using a Qubit RNA High Sensitivity Assay Kit (Thermo Fisher Scientific #Q32852) and an Agilent 2100 Bioanalyzer (Agilent Technologies, Santa Clara, CA, USA), respectively. The RNA eluates containing all fractions >200 nts were then aliquoted and stored at – 80°C prior to further processing.

### Sample Preparation for Nucleoside LC-MS/MS

A stable isotope-labelled internal standard (SILIS) was prepared as previously described (25). Briefly, *S. cerevisiae* was cultivated overnight (30°C, 250 rpm) in ^13^C,^15^N Silantes rich growth medium with 1% ^13^C_6_-glucose (Silantes, Bavaria, Germany). The culture was diluted to OD 0.1 with fresh medium (as above) to 100 mL and cultivated for two more days. The supernatant was removed by centrifugation (3000 xg, 5 min, 4°C) and the cells were resuspended in ultrapure water, transferred to clean tubes and pelleted again (3000 x g, 5 min, 4°C). The supernatant was removed and pellet was resuspended in 4 mL TES buffer and 4 mL acidic phenol. The sample was incubated at 65°C for 1h and vortexed in 15 min intervals. The mixture was incubated on ice for 5 min. and phases are separated by centrifugation (3000 x g, 5 min, 4°C). The aqueous phase was transferred to clean tube, mixed with 4 mL of chloroform and incubated for 5 minutes. This step was repeated once before precipitating the sample with ammonium acetate and ethanol (70%). The RNA was pelleted by centrifugation (12,000 g, 1h, 4°C) and resuspended in 250 µL ultrapure water.

Four samples of 10 µg of small RNA-depleted B-cell RNA in 20 µL of ultrapure water and 10 µg of SILIS were digested into single nucleosides by adding 16.5 µL of freshly prepared digestion mix. The final solution contained 2 U Benzonase, 2 U Alkaline Phosphatase (CIP), 0.2 U Phosphodiesterase I (PDE I), 1 µg Pentostatin, 5 µg Tetrahydrouridine and 10 pmol antioxidant butylated hydroxytoluene in a 5 mM Tris (pH 8), 1 mM MgCl2 buffer. After incubation at 37 °C for two hours, the samples were pooled and diluted with ultra-pure water to a concentration of 0.16 µg/µL. Two 110 µL aliquots were taken. To one of these, 11 µL of SILIS were added and to the other, 11 µL of ultra-pure water. For each instrument, 10 µL of both the SILIS-free and the SILIS-containing digested RNA were injected in triplicate.

Calibration solutions were prepared using synthetic nucleosides for absolute quantitative analysis and to define the linear range. Twelve calibration levels were created by serial dilution (with level 1 as the lowest and level 12 as the highest). The concentrations of these levels ranged from 195 nM to 400 µM for canonical nucleosides, from 9.8 nM to 20 µM for pseudouridine (Ψ) and dihydrouridine (D), and from 2.4 nM to 5 µM for all other modified nucleosides. To determine the linear range using an internal standard, 150 µL of each calibration solution was transferred to separate tubes, to which 15 µL of SILIS was then added. Then, 10 µL of each calibration solution (with and without SILIS) were injected for each instrument. Levels 2, 6 and 11 were injected a second time for quality control purposes.

### Nucleoside LC-MS/MS

Nucleoside LC-MS/MS was performed in parallel using three different types of MS instruments: 1) QqQ, 2) qTOF and 3) Orbitrap. The instruments used come from three different manufacturers: 1) Agilent (Santa Clara, CA USA), 2) Bruker (Billerica, MA USA) and 3) Thermo Fisher Scientific (Waltham, MA USA).

For separation, each instrument was run with an ACQUITY UPLC HSS T3 column (Waters, Milford, USA): 100 Å, 1.8 µm, 2.1 mm × 150 mm at a temperature of 30 °C and a flow rate of 0.4 mL/min. Mobile phase A was a 5 mM aqueous ammonium acetate buffer solution adjusted to a pH of 5.3 with glacial acetic acid (65 µL/L). Mobile phase B consisted of 40% acetonitrile and 60% mobile phase A. The gradient started with an initial isocratic hold at 100% mobile phase A for 7.6 min, followed by a linear increase to 2% mobile phase B at 16 min and 5% mobile phase B at 26.5 min. The organic content was then increased to 25% mobile phase B at 29.5 min, 50% mobile phase B at 32.3 min, and 75% mobile phase B at 36.4 min. After a 0.2 min hold at 75% mobile phase B, the column was washed with 99% mobile phase B from 39.6 to 46.8 min. Finally, the system was re-equilibrated to 100% mobile phase A within 0.1 min and these conditions were maintained until the end of the 55 min run.

Mass spectrometric analysis was performed on all instruments in positive ion mode. Data were acquired using Neutral Loss Scan (NLS) mode for the QqQ, while the qTOF and the Orbitrap were operated in data-dependent acquisition (DDA) mode. Detailed acquisition parameters and metadata are provided in Supplementary Table S1.

### Data Analysis of nucleoside LC-MS/MS

The annotation and the integration of signals for each nucleoside and, where applicable, the corresponding SILIS, was carried out using the vendor software for both samples and calibration standards.

#### Relative quantification (QqQ, qTOF, Orbitrap)

For relative quantification of sample prepared without and with SILIS, the peak areas of the nucleosides and, where applicable, their respective SILIS were exported to Excel.

For samples without SILIS, the nucleoside peak area multiplied by 1000 was normalized to the sum of the areas of all canonical nucleosides. (Eq. 1)

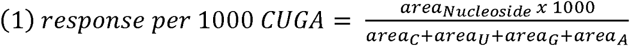

For the SILIS containing sample the ratio between the areas of the nucleoside and the corresponding SILIS was calculated to determine the Nucleoside Isotope Factor (NIF)(14) (Eq. 2):

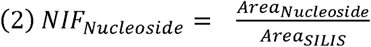

The 1000-fold NIF of each modified nucleoside was normalized to the sum of the NIFs of all canonical nucleosides (Eq. 3):

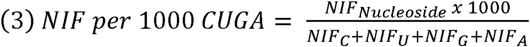

#### Determining linear range

To determine the linear range (both with and without SILIS), the peak areas of the nucleosides and, where applicable, the corresponding SILIS from the calibration standards were exported to Excel. For calibrations without SILIS, the nucleoside areas were normalized to the sum of the areas of all canonical nucleosides (Eq. 1). For calibrations with SILIS, the NIF was calculated first (Eq. 2) and then normalized to the sum of the NIFs of all canonicals (Eq. 3). The results for each calibration point were then related to the mean of all calibration points.

#### Absolute Quantification (qTOF, Orbitrap)

Absolute quantification was performed using R (26). The areas of the calibration standards were plotted against their respective molar amounts. The custom R script developed for this analysis is openly available via Zenodo (27). Calibration curves were created using linear, quadratic and power law fits with a weighting factor of 1/x², as recommended previously (28). This was done for the entire calibration range (calibration levels 1–12) and for a subset of the calibration range (calibration levels 6–11). The accuracy of the calibration levels and quality controls (QC) were calculated (Eq. 4), and the calibration curves were evaluated using the ICH criteria (Source: EMA/CHMP/ICH/172948/2019).

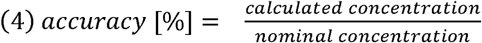

For further analysis, the simplest regression model that fit the calibration data over the range 6–11 was used: linear, quadratic or power law. Based on these curves, the molar amounts of nucleoside (n Nucleoside) in the samples were calculated. The molar amounts of each nucleoside were normalized to the sum of all canonicals (Eq. 5)

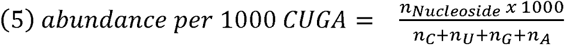

## Results

### Cross-platform study design and chromatography

We first established a unified experimental framework to assess retention time behavior across LC–MS platforms (Figure 1A). A defined mixture of 27 synthetic ribonucleoside standards, comprising both canonical and modified nucleosides, was used to determine retention times under controlled conditions, enabling a direct assessment of chromatographic behavior independent of biological variability. To isolate platform-dependent effects, chromatographic method settings were strictly matched across all measurements. All systems employed the same stationary phase, identical mobile phases prepared from a common stock, and matched gradient profiles. Sample preparation, injection volumes, and acquisition sequences were likewise aligned to minimize experimental variability. The three platforms represented distinct mass analyzer types (quadrupole, qTOF, and orbitrap-based detection) and ribonucleosides were identified by their precursor-ion, product-ion and retention time (individual injection of synthetic standards). Data processing was performed using the respective instrument specific software.

**Figure 1.**
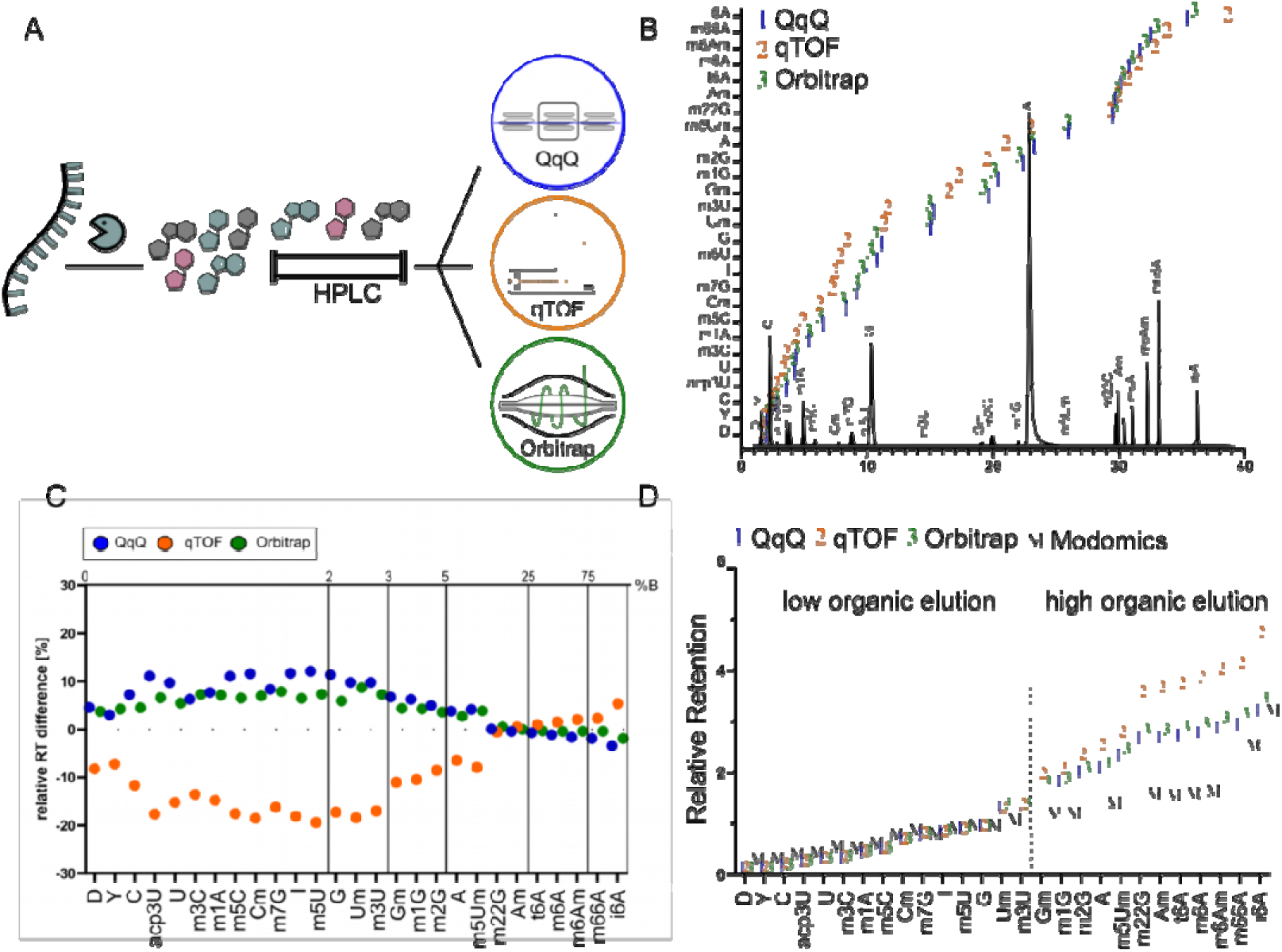
Study design and retention time behavior across LC–MS platforms. **(A)** Conceptual overview of the benchmarking workflow. A single, identical RNA sample was analyzed across three LC–MS platforms representing distinct mass analyzer types (QqQ, qTOF, Orbitrap) using fully harmonized chromatographic conditions (identical column, mobile phases, and gradients). Data were processed using vendor specific software to determine relative or absolute nucleoside amounts to then determine the extent of platform agreement. **(B)** Representative chromatogram illustrating the separation of canonical and modified ribonucleosides using a 55 minute gradient. Retention time (x-axis) is plotted against annotated nucleosides (y-axis). Data from all three platforms are overlaid (1, QqQ; 2, qTOF; 3, Orbitrap) (**C**) Relative retention time differences calculated from the mean retention time of all systems. Upper Y axis indicates gradient solvent composition with the grid indicating changes in solvent composition at the mean retention time of the corresponding modification. **(D)** Relative retention times normalized to guanosine for each platform (1, QqQ; 2, qTOF; 3, Orbitrap) and compared to MODOMICS reference values (M). Nucleosides are ordered by increasing retention.

As expected, all three LC–MS platforms produced consistent elution orders for canonical and modified nucleosides (Figure 1B). However, despite using the same column and programmed gradient, relative retention times calculated against the mean retention time across all systems differed between platforms (Figure 1C). Early-eluting nucleosides showed a higher relative difference than late-eluting nucleosides. This could be explained by differences in the design of the LC systems (e.g., varying tubing lengths from the multisampler to the column). These findings demonstrate that even when applying the exact same gradient to the same stationary phase do not guarantee identical retention times across LC–MS systems, highlighting a fundamental limitation for the transferability of retention-based annotations.

To facilitate comparison with the literature, we next calculated relative retention times (RT_rel_) by normalizing all nucleoside retention times to guanosine, a strategy widely adopted in the field and implemented in resources such as MODOMICS (1) (Figure 1D). This approach minimized the large retention time deviations observed for early-eluting nucleosides by compensating for differences in system dwell volume and flow path. However, systematic deviations remained for nucleosides eluting later in the chromatographic gradient (Figure S1B). As the gradient progresses, small differences in effective solvent composition accumulate between LC systems, resulting in retention shifts that cannot be corrected by normalization to a single reference compound. Although the use of multiple reference nucleosides distributed across the gradient could further improve retention time normalization (Figure S1C), such an approach would depend on the specific chromatographic method and is therefore difficult to generalize across laboratories. Consequently, relative retention times should be regarded as supportive rather than definitive identification criteria. Reliable nucleoside identification requires orthogonal evidence, including accurate mass, MS/MS (or MS³) fragmentation, and ideally comparison with authentic reference standards.

### Platform-dependent MS response behavior limits direct signal comparability

Total RNA from the B-cell line GM12878 was isolated and shipped from the US to Germany following the recently established shipping protocol of the human RNome project consortium (12). Chip gel electrophoresis revealed high integrity of the received RNA (Figure S2). At the same time, it was apparent that the sample mainly contained ribosomal RNA (rRNA) while small RNA such as tRNA was barely contained (Figure S1). The RNA was digested using a 1-pot digestion scheme (9,10) and split for analysis on the three platforms (Figure 2A-D). The QqQ platform was operated in neutral loss scan mode (NLS) (Figure 2A) as this mode best matches the DDA (data dependent acquisition) of the qTOF and orbitrap platforms (Figure 2B, C).

**Figure 2.**
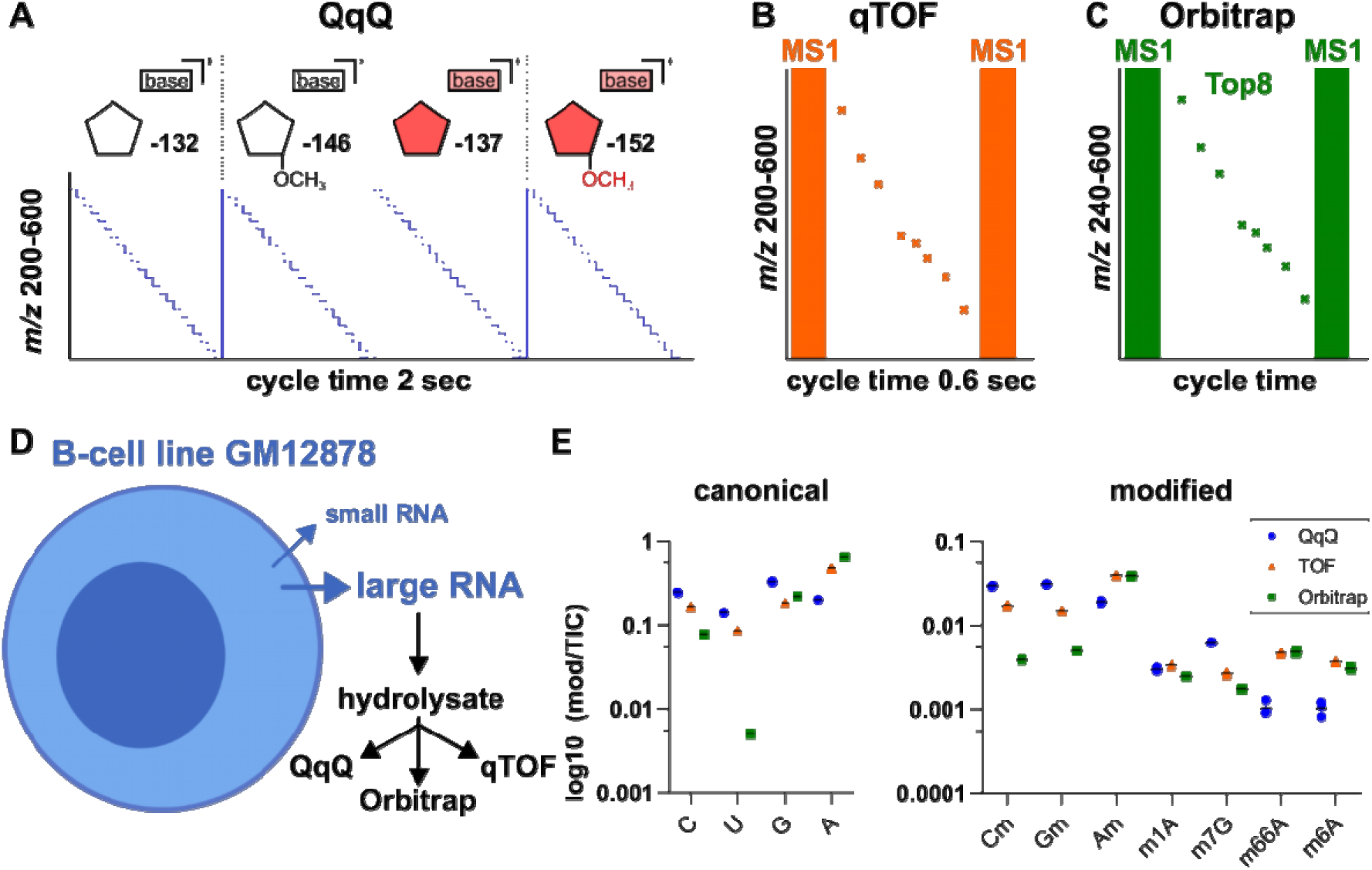
Acquisition-mode-specific detection of modified ribonucleosides. **(A)** Schematic representation of the NLS acquisition used on the QqQ system **(B)** Schematic representation of the DDA acquisition used on the qTOF system (fixed cycle time 0.6 sec) **(C)** Schematic representation of the DDA acquisition used on the orbitrap system (variable cycle time) **(D)** small-RNA depleted RNA from B-Cell GM12878 was hydrolyzed and 1.6 µg injected into each system. **(E)** relative MS signal of canonical (left) and modified (right) nucleosides measured across three LC–MS platforms under identical experimental conditions. Intensities are shown on a log10 scale. From n=3 replicates, all datapoints are shown, error bars reflect standard deviation.

Briefly, DDA is a detection mode scans for the highest abundant ions before selecting them for fragmentation and subsequent scanning of product ions. NLS is similar to DDA, as ions are scanned and fragmented before the resulting product ions are detected. The main difference is that the QqQ only detects ions that show a predefined mass difference. For nucleoside MS, this mass difference typically is 132 Da – the mass loss of ribose caused by fragmentation of the glycosidic bond. Though previously applied (29,30), NLS is not an ideal detection mode for QqQ platforms but was used due its similarly to DDA, the common acquisition mode for qTOF and orbitrap instruments. Commonly, QqQ instruments are used in multiple reaction monitoring (MRM) mode which was recently benchmarked by Hengesbach et al. (6) within the framework of the Human RNome project.

First data analysis from all platforms revealed that modifications associated with small RNAs were detectable on all three devices, but at low abundance. Thus, we focused on the abundant rRNA modifications for quantitative comparison of platforms and harmonization strategies. Despite identical sample composition and chromatographic conditions, signal intensities differed systematically between systems and spanned multiple orders of magnitude (Figure 2E). These differences were compound-specific and not explained by a simple global scaling factor, indicating platform-dependent response behavior for individual nucleosides. Importantly, observed signal differences do not reflect analytical sensitivity but rather system-specific ionization and detection characteristics. As a result, raw MS signal intensities are not directly comparable across platforms.

To assess the extent of platform dependent response differences, we first quantified selected modifications in identical RNA samples using normalized MS signal intensities (Figure 3A). For this analysis, the MS signal of each modification was normalized to the RNA input (MS signal of 10³ canonical nucleosides). The resulting relative modification levels differed substantially between platforms. The observed deviations were analyte dependent and varied in both magnitude and direction, demonstrating that quantification is strongly influenced by compound-specific response behavior rather than by uniform platform-specific scaling effects. Furthermore, even on the same platform, relative quantities varied substantially across different analyte amounts, due to deviation from linear behavior over large parts of the concentration range (Figure 3B). As a result, the relationship between modification and canonical signals changed depending on the injected amount, thereby directly affecting quantification. Together, these results demonstrate the poor inter-platform agreement and show that relative MS signal ratios alone cannot provide harmonized quantification.

**Figure 3.**
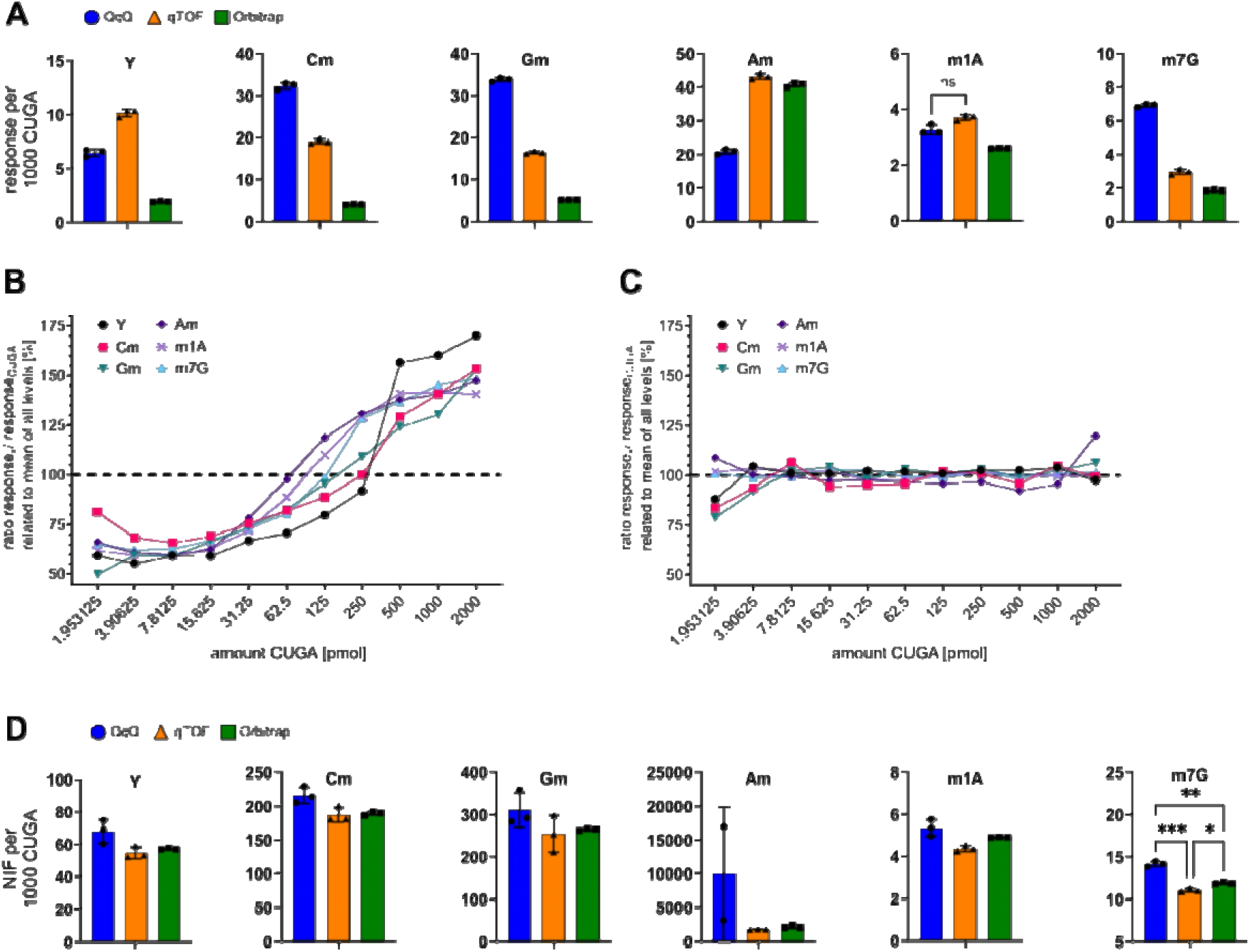
SILIS correction enables comparable relative quantification across platforms by correcting compound-specific response behavior. **(A)** Relative quantification of selected RNA modifications calculated as the ratio of modification signal to 10^3^ nucleosides (1000 CUGA) (n = 3; error bars, s.d., (Brown-Forsythe/Welch ANOVA, followed by Dunnett‣s T3 post-hoc test for multiple comparison, P < 0.05 for all comparisons; not indicated for clarity)). **(B)** Concentration-dependent response behavior of modification-to-canonical signal ratios without internal standard correction. **(C)** Concentration-dependent response behavior of modification-to-canonical signal ratios with internal standard correction. **(D)** SILIS-based quantification expressed as the ratio of normalized modification and canonical signals (NIF=Nucleoside-Isotope-Factor (29)) (n = 3; error bars, s.d., with no statistically significant differences between systems unless indicated (Brown-Forsythe/Welch ANOVA, followed by Dunnett‣s T3 post-hoc test for multiple comparison, * = P ≥ 0.05; ** = P ≥ 0.01; *** = P ≥ 0.001)).

### SILIS correction enables harmonized relative quantification across LC–MS platforms

We next repeated relative quantification using stable isotope-labeled internal standards (SILIS) (25,29). Even within the same platform, SILIS correction stabilized the relative response, over a large concentration range by substantially extending the linear dynamic range (Figure 3C). In contrast to uncorrected relative quantification (Figure 3A), SILIS-corrected values showed strong agreement between platforms, with substantially reduced inter-system variability across all analyzed modifications (Figure 3D). As a result, quantitative values showed markedly improved comparability across the evaluated workflows. Still, residual analyte specific differences remained and in case of m^7^G are even statistically significant. A plausible explanation for this deviation is the frequent co-elution of m^7^G with abundant digestion additives utilized in our assay, such as tetrahydrouridine (THU) or butylated hydroxytoluene (BHT). While SILIS effectively compensates for moderate ion suppression, severe matrix effects caused by such co-eluting compounds can exceed the corrective capacity of the internal standard. Furthermore, the addition of SILIS can introduce cumulating variances. For instance, the quantification of Am on the Agilent platform exhibited large standard deviations only after SILIS correction, whereas the uncorrected values showed low variance, mainly because too few datapoints were generated for the SILIS peak, reducing precision and accuracy of the calculated ratio. This highlights that while internal standardization generally improves cross-platform comparability, it does not completely rescue severe matrix effects and, under certain conditions, can contribute to signal variance.

### Calibration range selection determines accuracy of absolute quantification

For absolute quantification, freshly prepared stock solutions of synthetic nucleosides were used (17). The present benchmark focused on identifying appropriate calibration strategies for cross-platform comparison rather than performing complete bioanalytical validation of each individual workflow according to regulatory guidelines. Consequently, parameters such as formal Lower Limit of Quantification (LLOQ) or Upper Limit of Quantification (ULOQ) determination, recovery experiments, or long-term precision were not established for every analyte and platform.

To establish a calibration strategy for absolute quantification, we evaluated the regression models according to ICH guidelines. Model performance was assessed based on the back-calculated accuracy of the calibration levels and quality controls (QCs). Specifically, at least 75% of the non-zero calibration standards must fall within ±15% (85–115%) of their nominal concentration, except at the lowest calibration point, where a deviation of ±20% (80–120%) is acceptable (ICH M10) (31). The simplest regression model fulfilling these criteria was then selected for quantification. Ideally, the optimal weighting factor would be determined empirically via variance analysis. However, due to the time constraints inherent to the workshop format calibrations were measured in unicate, making this approach unfeasible. Therefore, 1/x^2^ was selected for all calibration curves, representing a standard weighting factor accepted for LC–MS/MS applications (28).

Initially, we applied this approach to evaluate the performance of calibration models across an extended concentration range using 12 calibration levels. In general, fitting is straightforward within the linear range, while extending the range towards detector saturation leads to clear deviations from linearity observed at higher concentrations (Figure 4A). Although quadratic or power-law models can extend the quantifiable range, expanding the concentration range too far eventually leads deviations that cannot be adequately described by any acceptable regression model. Employing more complex models at this stage would introduce a high risk of overfitting. To address this issue, the calibration range was reduced to six concentration levels, that covered the concentration of our sample. Under these conditions, regression models showed markedly improved consistency across analytes, in line with established ICH recommendations for quantitative LC–MS workflows (Figure 4A). Linear regression provided an adequate fit in most cases, whereas a subset of analytes required quadratic regression to account for residual curvature. Only rarely, a power law model was necessary, indicating pronounced non-linearity that could not be captured by polynomial fitting. A systematic comparison of regression models across all detected nucleoside modifications confirmed this trend (Figure 4B). These results demonstrate that rather than forcing a fit over an unnecessarily wide range, restricting the calibration to a well-defined concentration range around sample concentration enables more accurate regression modelling, forming the basis for subsequent absolute quantification. Following optimization of the calibration strategy, we performed absolute quantification of selected RNA modifications in the native sample using the established 6-point calibration models (Figure 4D). The selected regression models, along with information on the specific QCs and calibration levels that met the acceptance criteria, are summarized in (Table S2). Absolute abundances were calculated for each modification based on external calibration and normalized to total nucleoside content, expressed as amount per 10^3^ canonicals. Quantification was carried out on qTOF and orbitrap platforms, as the QqQ system did not provide sufficient data density across chromatographic peaks due to its acquisition cycle time, precluding reliable peak integration. Importantly, external calibration substantially improved cross-platform agreement in comparison to direct relative quantification of raw signals. However, without internal standard correction, residual variability between systems remained evident for several analytes. (Figure 4C) This indicates that external calibration alone is insufficient to achieve complete cross-platform agreement. Together, these results demonstrate that absolute quantification based on controlled external calibration enables more consistent determination of RNA modification quantities but failed to eliminate residual variability between systems. Evaluation of SILIS-based absolute quantification was beyond the scope of this study due to constraints from workshop structure and timeline.

**Figure 4.**
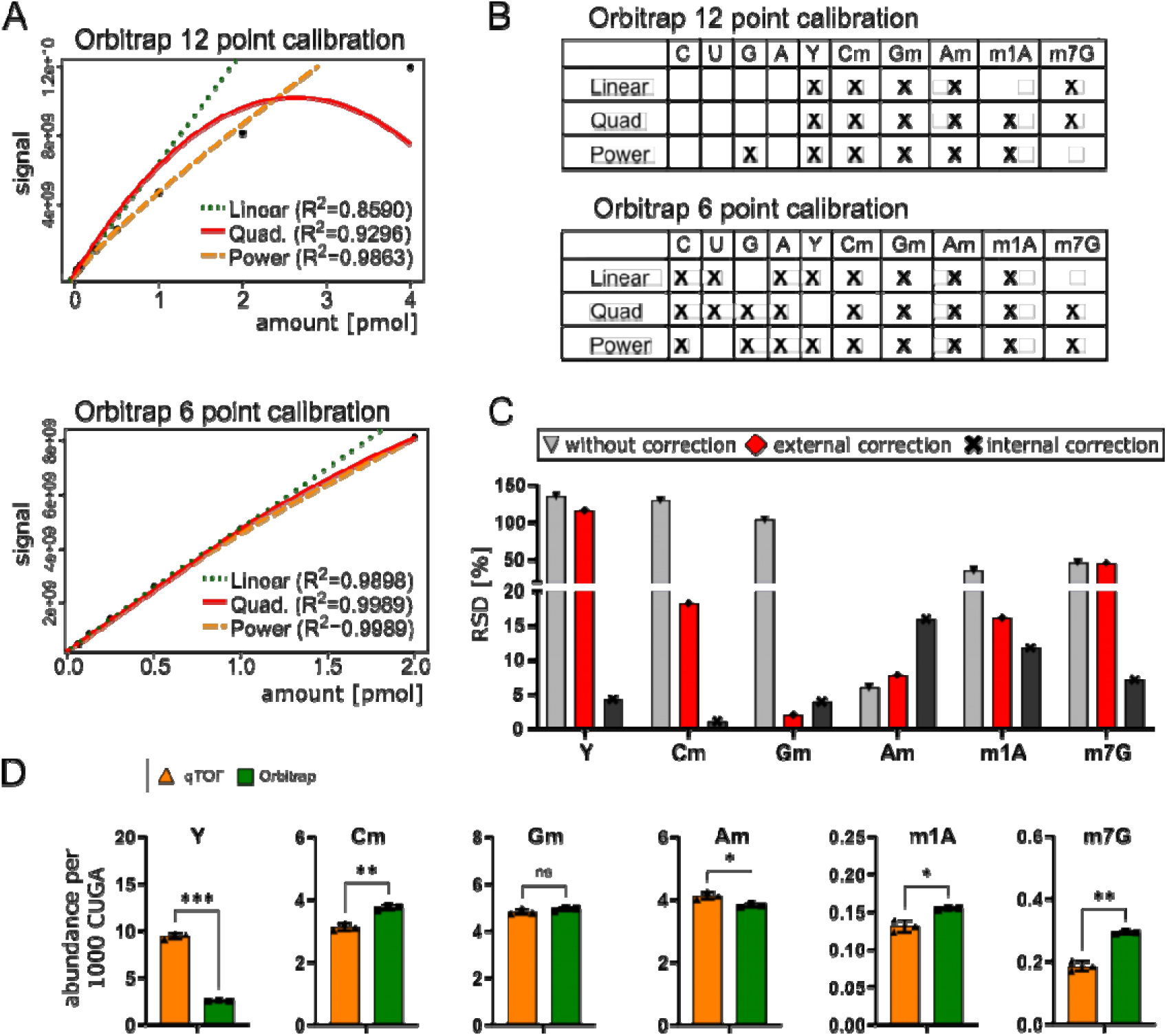
Calibration strategy and absolute quantification of RNA modifications across LC–MS platforms. **(A)** Representative calibration curves generated using 12-point and 6-point calibration series across an extended concentration range. **(B)** Distribution of regression models required to adequately describe calibration behavior across all detected nucleosides. (**C**) Relative percent difference (RPD) between qTOF and Orbitrap for relative quantification without SILIS (without correction), relative quantification with SILIS (internal correction) and absolute quantification (external correction) **(D)** Absolute quantification of selected RNA modifications in the native sample based on external calibration and normalized to total nucleoside content (per 1,000 nucleotides, CUGA). Quantification was performed on qTOF (orange) and Orbitrap (green) platforms. (Welch t-test, * = P < 0.05; ** = P < 0.01; *** = P < 0.001).

### Benchmark-derived principles for robust nucleoside LC–MS/MS workflows

LC–MS-based RNA modification analysis encompasses analytically distinct objectives ranging from qualitative structure validation to relative and absolute quantification (Figure 5). While all objectives face equal challenges at the stage of sample preparation, digestion and general LC-MS/MS conditions, each downstream analytical goal requires measures to ensure data integrity. The complete workflow and best practices, identified from this study and recent literature (9,10,17,20), are summarized in Figure 5. In the following section we will briefly discuss the state-of-the-art for each part of the workflow.

**Figure 5.**
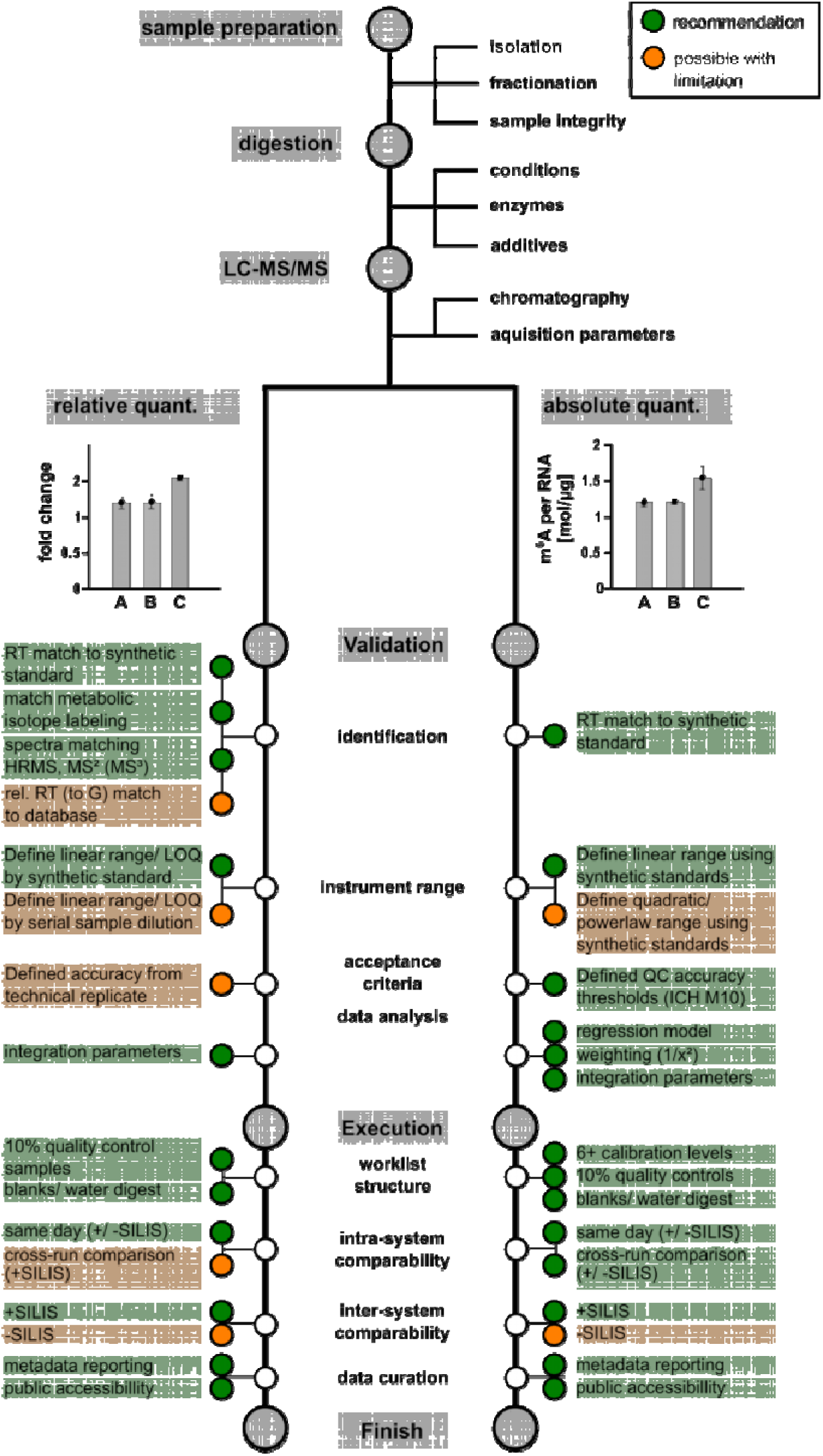
Analytical framework for qualitative and quantitative nucleoside MS workflows. The workflow is divided into common upstream analytical steps including sample preparation, RNA digestion, and LC–MS/MS acquisition, followed by two downstream analytical objectives: relative quantification (normalization of MS response), and absolute quantification. Green labels indicate parameters considered recommended for robust quantification. Orange labels indicate context-dependent or partially sufficient parameters. Shared nodes indicate workflow components or validation steps that contribute to multiple analytical objectives. The framework highlights that increasing quantitative transferability requires progressively stricter standardization of calibration, normalization, and data analysis strategies across laboratories and mass spectrometry platforms.

### Sample preparation and RNA integrity

Sample preparation is a major determinant of reproducible nucleoside MS analysis and directly affects all downstream identification and quantification steps. Since nucleoside mass spectrometry depends entirely on the quality of the RNA input, differences in isolation strategy, RNA fractionation, or sample purity directly impact the biological relevance of the results. For example, TRIzol- and kit-based workflows may differ with respect to salt carryover, residual contaminants, and recovery of specific RNA classes. Likewise, RNA fractionation strongly influences the observed modification profile, as RNA modifications are highly RNA-class dependent. A prominent example is the purification of mRNA for analysis of m^6^A abundance. Here, residual rRNA impacts the result and stringent removal is required (32,33). Similarly, rRNA fragments may contaminate small RNA fractions which impacts quantification of tRNA modifications (34). Consistent separation workflows and careful annotation of the analyzed RNA population are therefore essential. In addition, RNA integrity and sample purity must be rigorously controlled prior to LC–MS analysis. We therefore recommend systematic quality control using electrophoretic integrity analysis (e.g., chip electrophoresis) prior to nucleoside analysis.

### Digestion-associated artifacts

Enzymatic digestion represents another critical source of variability in nucleoside LC–MS workflows and can directly influence both quantitative accuracy and structure analysis of modified nucleosides. Chemical reactivity of nucleosides (10,18,21) and enzymatic reactivity can lead to misidentification (20) or wrong quantitative assessment. The choice of nuclease cocktails for digestion must be carefully selected depending on which modifications are of the main interest of the analysis (10). Furthermore, Nucleases (and probably many other enzymes) are contaminated with deaminases that can trigger artificial A-to-I or C-to-U conversion in DNA (35) and RNA (10). Therefore, inclusion of deaminase inhibitors (coformycine/pentostatin and tetrahydrouridine) must be evaluated (9). **Chromatographic and ionization effects.** Chromatographic separation critically influences ionization efficiency and quantitative accuracy in LC–MS/MS-based nucleoside analysis. Matrix-derived ion suppression, and partial co-elution of other nucleosides or additives (note: coformycine/pentostatine may co-elute with G, depending on the selected gradient) may affect signal intensities and thereby introduce potential biases into quantitative measurements (10). Crucially, when methods are transferred between platforms, inter-system hardware differences, such as varying mixer sizes or post-injector dwell volumes, cause subtle shifts in exact retention times. As a result, the same analyte may elute at a slightly different effective solvent composition depending on the system. This variation in the mobile phase reaching the ion source directly alters electrospray ionization (ESI) efficiency. These effects are especially relevant in workflows lacking SILIS (25,29), where LC-MS variability directly propagates into quantitative uncertainty.

### MS Acquisition

MS acquisition strategy further contributes to variability in quantitative performance. Data-dependent acquisition (DDA) and neutral loss scanning (NLS) enable broader modification discovery and increase analytical depth, but also introduce additional complexity that can compromise quantitative robustness, particularly for low abundant analytes or partially coeluting species. Therefore, targeted analysis has become the field standard for quantitative approaches (9,25,29,34). Targeted methods likewise require careful optimization of acquisition parameters to guarantee high quality signal for all analytes of interest. Source, transmission, fragmentation and detector parameters must be optimized to ensure that signal for all analytes of interest surpasses threshold signal-to noise ratio necessary for quantification. Additionally, instrument cycle time must be adjusted to fit the expected chromatographic peak width to ensure adequate data depth (minimum 10-15 point per peak) (36), effectively limiting the number of transitions that can be reliably monitored within the same time window.

### Qualitative / Structural analysis

In both relative and absolute quantification regimes, identification of the nucleosides of interest within the chromatographic gradient is a prerequisite for quantitative analysis.

This can be complicated in practice by isobaric modifications, positional isomers and naturally occurring isotopologues whose overlapping MS signals lead to wrong signal assignment. For unambiguous identification at least three parameters of a nucleoside must match the expectation: precursor ion *m/z* as determined in a full MS spectrum, MS/MS spectrum fragmentation match and retention time must fit synthetic standard. If a synthetic standard is not available, RT_rel_ values from MODOMICS (against G) may be an alternative to ensure consistency with expected elution order. Furthermore, fragmentation of the base (MS³) may be used and cross-referenced with the literature (37–40) to ensure a nucleoside’s identity. Finally, metabolic labelling has supported the validation of modified nucleoside structures in the past (5,30,41–43). A detailed guideline to nucleoside identification can also be found here (20).

### Quantitative analysis

While both LC–MS workflows share common upstream analytical challenges during sample preparation, digestion, and data acquisition, the downstream analytical requirements differ substantially depending on the intended quantitative objective. Relative and absolute quantification require differing levels of analytical validation, normalization control, and inter-system harmonization.

### Relative Quantification

The most common objective in quantitative nucleoside MS is assessment of relative differences in modification content between biological conditions e.g. treatment, stress conditions or different RNA types. Typically, some form of normalization is required to control for differences in input material quantity, so modification signal areas are related to those of one or more canonical nucleosides or RNA amount estimated by other methods (e.g. UV quantification or chip gel electrophoresis). The obtained ratios are numerically arbitrary and do not reflect modification stoichiometry but can be used to calculate fold changes between conditions. Relative quantification is comparatively robust as instrument specific response differences cancel out for intra-system comparisons. Crucially, measurement in linear range is a prerequisite to obtain meaningful results, therefore dynamic ranges must be validated for both the modification of interest and for the canonical nucleosides used for normalization. For this, synthetic standard or serial dilution of biological sample can be used. Differences in ionization efficiency between batches cannot be excluded so application of internal standards is still recommended. As we could demonstrate, the use of SILIS compensates for ionization differences and thereby increases the linear range (Figure 3 B/C). For inter-system comparison of relative quantitative data from this study, use of an internal standard was necessary and sufficient to ensure comparability between LC-MS platforms of fundamentally different architecture (Figure 3D). This is in accordance with previous observations of SILIS correction reducing instrument variability across time (14). We focused on comparatively abundant modifications quantified from identical large RNA pools, so potential limitations for transferability to low abundant species and other sample types should be noted.

Furthermore, for both intra-, and intersystem comparisons, worklist structure must support integrity of acquired data by inclusion of blank injections, water digest to control for matrix background and quantitative quality control samples to verify consistent instrument response. Although there are no official recommendations for peak integration tools (e.g. data smoothing, integration algorithms, and baseline treatment), it is fundamental that all samples are treated consistently.

**Absolute quantification** determines the abundance of modified ribonucleosides in defined physical units, such as fmol per µg RNA or modifications per cell, and is required for stoichiometric analysis, inter-study comparison, biomarker development, and quantitative systems biology approaches. In contrast to relative quantification, absolute quantification requires experimentally validated calibration models capable of converting instrument response into absolute analyte amounts. This introduces additional analytical requirements, including definition of calibration ranges, regression model validation, weighting strategies, quality-control criteria, and determination of signal inclusion criteria.

As demonstrated in this benchmark, excessively broad calibration ranges frequently impaired regression accuracy, whereas restricted calibration ranges enabled more reproducible model fitting across analytes and platforms. We therefore recommend experimentally defining instrument and analyte specific calibration ranges together with systematic evaluation of regression model and quality-control performance prior to quantitative analysis in accordance with ICH M10 guidelines (31). Regression validation should comprise weighting of obtained regression curves and should be applied consistently across runs. The correct weighting factor can be identified by analysis of variance across calibration levels (28). In terms of integration the same recommendations laid out above apply. For worklist structure a minimum of six calibration points must be considered.

## Discussion

The present benchmark identifies core limitations in the comparability of relative and absolute nucleoside MS workflows across instrument architectures. While chromatographic reproducibility was high across platforms, quantitative agreement varied across analytes and correction method. These findings highlight that reproducible epitranscriptomic analysis cannot be achieved through isolated optimization of individual workflow components alone but instead requires harmonized control of the complete analytical process.

It is important to note that executing strictly matched LC methods on matched columns does not guarantee identical chromatography across different instruments. Intersystem differences, such as varying dead volumes and flow path length, inevitably lead to subtle variations in retention times and gradient delivery, making absolute chromatographic behavior inherently platform-dependent. Comparison of relative retention times partially improves agreement across instruments. Crucially, this normalization is only effective within a defined gradient section (Figure S1), therefore transferability across different systems and gradients is limited. Differences in elution time translate into differences of ionization efficiency, thereby highlighting the need for analytical standards for quantitative approaches.

External calibration improved comparability between systems but did not fully eliminate platform-dependent response behavior, indicating that external calibration alone is insufficient to ensure transferable absolute quantification. Future cross-platform benchmarking studies should therefore implement harmonized correction architectures comprising centrally distributed calibration standards, stable isotope-labeled internal standards, validated normalization strategies, and predefined reporting criteria. In parallel, analyte-specific validation datasets should be generated, including calibration residuals, back-calculated calibrator accuracy, quality-control precision, variance-based weighting factor selection, and determination of the applicable calibration range. Such multicenter validation studies would establish the analytical foundation required to define transferable quantitative performance criteria for nucleoside LC–MS and ultimately enable robust absolute quantification across laboratories and mass spectrometry platforms.

Importantly, internal correction strategies such as SILIS correction cannot compensate for fundamentally unsuitable analytical conditions. SILIS substantially improved relative quantification by correcting compound-specific response behavior across platforms and concentrations. However, isotope dilution does not rescue insufficient chromatographic separation, inappropriate acquisition settings, detector saturation, or inadequate signal sampling across chromatographic peaks. Retrospective data analysis showed that on the QqQ platform, suboptimal dwell time and cycle time settings were chosen and resulted in insufficient data point density across chromatographic peaks, thereby substantially limiting quantitative accuracy despite otherwise harmonized chromatographic conditions. These limitations are a consequence of scan-mode acquisition, and are generally less problematic for targeted measurements but still highlight dwell and cycle time as sensitive parameters that must be considered to achieve robust quantitative nucleoside MS.

We further observed that downstream data analysis is not a neutral processing step but instead directly contributes to analytical variability. Peak detection, smoothing, integration boundaries, and signal filtering all influence both modification identification and quantitative output. As a result, data processing workflows themselves represent part of the analytical measurement architecture and require harmonized implementation and transparent reporting to ensure interlaboratory reproducibility. This aspect is particularly relevant for modifications of low-abundance and partially co-eluting signals, where small differences in peak integration strategy may substantially alter quantitative interpretation.

Importantly, several instrument platforms offer targeted acquisition modes specifically designed for quantitative analysis, but such platform-specific optimizations were intentionally not implemented in the present benchmark in order to maintain a harmonized acquisition framework across systems. In particular, the qTOF and orbitrap systems employed DDA-based acquisition strategies that are ideally suited for qualitative structure analysis and modification discovery but are less optimized for quantitative analysis. Likewise, the extended chromatographic gradient used in this study was intentionally selected to minimize co-elution and ion suppression effects rather than maximize throughput. While modern targeted nucleoside workflows increasingly employ substantially shorter gradients, often in the range of 5–15 minutes (25,44), the present study prioritized analytical robustness and cross-platform comparability under heterogeneous real-world conditions. We therefore expect that the harmonization principles identified here will be directly transferable, and potentially further improved, in dedicated targeted quantitative workflows employing optimized acquisition methods.

## Future directions toward transferable interlaboratory RNA modification quantification

The present benchmark demonstrates that harmonized chromatography and shared analytical workflows alone are insufficient to ensure transferable relative and absolute nucleoside quantification across laboratories and mass spectrometry platforms. Instead, future benchmarking efforts must move toward standardized correction architectures that systematically address normalization, calibration transferability, and platform-dependent response effects.

A next-generation benchmarking framework should therefore include centrally distributed reference materials with defined and experimentally validated modification stoichiometries. In particular, synthetic RNAs containing known modification patterns and systematically varied ratios of canonical to modified nucleosides would enable controlled assessment of normalization strategies across concentration ranges and analytical platforms. Such materials could serve as universal validation standards for future epitranscriptomic workflows and provide a defined basis for interlaboratory comparison.

In parallel, systematic dilution series should be incorporated into future benchmark designs to experimentally determine under which normalization strategies equivalent quantitative results can be achieved across platforms and concentration ranges. This will be essential to define the limits of transferable quantification and to distinguish robust correction strategies from normalization approaches that remain platform-dependent. The benchmark further highlights the need for centrally available stable isotope-labeled internal standard mixtures (SILIS) with defined composition and harmonized spike-in procedures. Likewise, calibration solutions should be centrally prepared and distributed to participating laboratories to minimize variability introduced by independent preparation, storage, or concentration assignment. Together, such standardized reference systems would enable rigorous evaluation of inter-system comparability and establish the analytical foundation required for reproducible large-scale epitranscriptomic studies across laboratories and instrument platforms.

## Data availability

The data supporting this article have been included as part of the Supplementary Information.

## Acknowledgements

This work originated from the 2025 Human RNome Project Consortium (HRPC) Workshop held in Frankfurt, Germany. The workshop was made possible through the generous support of the Warren Alpert Foundation, Brown University, Goethe University Frankfurt, Vereinigung der Freunde und Förderer der Goethe-Universität, the Deutsche Pharmazeutische Gesellschaft (DPhG), House of Pharma and Healthcare, the Collaborative Research Centre SFB 1309, the RNA Modification and Processing (RMaP) initiative, Bruker, Thermo Fisher Scientific, Alida Biosciences, and Biospring. Their support enabled the international gathering of the consortium and the collaborative benchmarking study presented here. The authors further thank all workshop participants for their scientific contributions, open discussions, and collaborative spirit, which laid the foundation for this consortium study. We gratefully acknowledge time and resources invested by our industry partners Thermo Fisher Scientific and Bruker Daltonics GmbH & Co. KG in support of the workshop. We thank Birgit Bartussek for her administrative and organizational contribution to the workshop and all former and current members of the Kaiser/Kellner lab.

## References

1. Cappannini, A., Ray, A., Purta, E., Mukherjee, S., Boccaletto, P., Moafinejad, S.N., Lechner, A., Barchet, C., Klaholz, B.P., Stefaniak, F. et al. (2024) MODOMICS: a database of RNA modifications and related information. 2023 update. Nucleic Acids Res, 52, D239–D244,doi: 10.1093/nar/gkad1083.

2. Bessler, L., Vogt, L.M., Lander, M., Dal Magro, C., Keller, P., Kuhlborn, J., Kampf, C.J., Opatz, T. and Helm, M. (2023) A New Bacterial Adenosine-Derived Nucleoside as an Example of RNA Modification Damage. Angew Chem Int Ed Engl, 62, e202217128,doi: 10.1002/anie.202217128.

3. Kimura, S., Dedon, P.C. and Waldor, M.K. (2020) Comparative tRNA sequencing and RNA mass spectrometry for surveying tRNA modifications. Nat Chem Biol, 16, 964–972,doi: 10.1038/s41589-020-0558-1.

4. Miyauchi, K., Kimura, S., Akiyama, N., Inoue, K., Ishiguro, K., Vu, T.S., Srisuknimit, V., Koyama, K., Hayashi, G., Soma, A. et al. (2025) A tRNA modification with aminovaleramide facilitates AUA decoding in protein synthesis. Nat Chem Biol, 21, 522–531,doi: 10.1038/s41589-024-01726-x.

5. Reichle, V.F., Petrov, D.P., Weber, V., Jung, K. and Kellner, S. (2019) NAIL-MS reveals the repair of 2-methylthiocytidine by AlkB in E. coli. Nat Commun, 10, 5600,doi: 10.1038/s41467-019-13565-9.

6. Gatsiou, A. and Stellos, K. (2023) RNA modifications in cardiovascular health and disease. Nat Rev Cardiol, 20, 325–346,doi: 10.1038/s41569-022-00804-8.

7. Chujo, T. and Tomizawa, K. (2025) Neurological Diseases Caused by Loss of Transfer RNA Modifications: Commonalities in Their Molecular Pathogenesis. J Mol Biol, 437, 169047,doi: 10.1016/j.jmb.2025.169047.

8. Watts, J.A., Grunseich, C., Rodriguez, Y., Liu, Y., Li, D., Burdick, J.T., Bruzel, A., Crouch, R.J., Mahley, R.W., Wilson, S.H. et al. (2022) A common transcriptional mechanism involving R-loop and RNA abasic site regulates an enhancer RNA of APOE. Nucleic Acids Res, 50, 12497–12514,doi: 10.1093/nar/gkac1107.

9. Cai, W.M., Chionh, Y.H., Hia, F., Gu, C., Kellner, S., McBee, M.E., Ng, C.S., Pang, Y.L., Prestwich, E.G., Lim, K.S. et al. (2015) A Platform for Discovery and Quantification of Modified Ribonucleosides in RNA: Application to Stress-Induced Reprogramming of tRNA Modifications. Methods Enzymol, 560, 29–71,doi: 10.1016/bs.mie.2015.03.004.

10. Ammann, G., Berg, M., Dalwigk, J.F. and Kaiser, S.M. (2023) Pitfalls in RNA Modification Quantification Using Nucleoside Mass Spectrometry. Acc Chem Res, 56, 3121–3131,doi: 10.1021/acs.accounts.3c00402.

11. Basanta-Sanchez, M., Temple, S., Ansari, S.A., D’Amico, A. and Agris, P.F. (2016) Attomole quantification and global profile of RNA modifications: Epitranscriptome of human neural stem cells. Nucleic Acids Res, 44, e26,doi: 10.1093/nar/gkv971.

12. Hengesbach, M., Chan, C.K., Bhandari, T., Bruzel, A., DeMott, M.S., Podoprygorina, G., Sun, G., Tabeling, E., Cheung, V.G., Dedon, P.C. et al. (2025) Toward standardized epitranscriptome analytics: an inter-laboratory comparison of mass spectrometric detection and quantification of modified ribonucleosides in human RNA. Nucleic Acids Res, 53,doi: 10.1093/nar/gkaf895.

13. Sarin, L.P., Kienast, S.D., Leufken, J., Ross, R.L., Dziergowska, A., Debiec, K., Sochacka, E., Limbach, P.A., Fufezan, C., Drexler, H.C.A. et al. (2018) Nano LC-MS using capillary columns enables accurate quantification of modified ribonucleosides at low femtomol levels. RNA, 24, 1403–1417,doi: 10.1261/rna.065482.117.

14. Kellner, S., Ochel, A., Thuring, K., Spenkuch, F., Neumann, J., Sharma, S., Entian, K.D., Schneider, D. and Helm, M. (2014) Absolute and relative quantification of RNA modifications via biosynthetic isotopomers. Nucleic Acids Res, 42, e142,doi: 10.1093/nar/gku733.

15. He, L., Wei, X., Ma, X., Yin, X., Song, M., Donninger, H., Yaddanapudi, K., McClain, C.J. and Zhang, X. (2019) Simultaneous Quantification of Nucleosides and Nucleotides from Biological Samples. J Am Soc Mass Spectrom, 30, 987–1000,doi: 10.1007/s13361-019-02140-7.

16. Sun, J., Wu, J., Yuan, Y., Fan, L., Chua, W.L.P., Ling, Y.H.S., Balamkundu, S., Dwijapriya, Suen Suen H.C., de Crécy-Lagard, V., et al. (2024) tRNA modification profiling reveals epitranscriptome regulatory networks in *Pseudomonas aeruginosa*. *bioRxiv*, 10.1101/2024.07.01.601603, 2024.2007.2001.601603,doi: 10.1101/2024.07.01.601603.

17. Kerkhoff, K., Wesseling, H., Qi, Y., Obersteiner, S., Liu, K., Berg, M., Rusling, L., Zipse, H. and Kaiser, S. (2026) Long-term stability of RNA nucleoside standards for accurate LC-MS quantification. Nucleic Acids Res, 54,doi: 10.1093/nar/gkag444.

18. Jora, M., Borland, K., Abernathy, S., Zhao, R., Kelley, M., Kellner, S., Addepalli, B. and Limbach, P.A. (2021) Chemical Amination/Imination of Carbonothiolated Nucleosides During RNA Hydrolysis. Angew Chem Int Ed Engl, 60, 3961–3966,doi: 10.1002/anie.202010793.

19. Bessler, L., Gross, J., Kampf, C.J., Opatz, T. and Helm, M. (2024) Reversible oxidative dimerization of 4-thiouridines in tRNA isolates. RSC Chem Biol, 5, 216–224,doi: 10.1039/d3cb00221g.

20. Kaiser, S., Byrne, S.R., Ammann, G., Asadi Atoi, P., Borland, K., Brecheisen, R., DeMott, M.S., Gehrke, T., Hagelskamp, F., Heiss, M. et al. (2021) Strategies to Avoid Artifacts in Mass Spectrometry-Based Epitranscriptome Analyses. Angew Chem Int Ed Engl, 60, 23885–23893,doi: 10.1002/anie.202106215.

21. Matuszewski, M., Wojciechowski, J., Miyauchi, K., Gdaniec, Z., Wolf, W.M., Suzuki, T. and Sochacka, E. (2017) A hydantoin isoform of cyclic N6-threonylcarbamoyladenosine (ct6A) is present in tRNAs. Nucleic Acids Res, 45, 2137–2149,doi: 10.1093/nar/gkw1189.

22. Espadas, G., Morales-Sanfrutos, J., Medina, R., Lucas, M.C., Novoa, E.M. and Sabido, E. (2022) High-performance nano-flow liquid chromatography column combined with high- and low-collision energy data-independent acquisition enables targeted and discovery identification of modified ribonucleotides by mass spectrometry. J Chromatogr A, 1665, 462803,doi: 10.1016/j.chroma.2022.462803.

23. Ross, R.L., Yu, N., Zhao, R., Wood, A. and Limbach, P.A. (2023) Automated Identification of Modified Nucleosides during HRAM-LC-MS/MS using a Metabolomics ID Workflow with Neutral Loss Detection. J Am Soc Mass Spectrom, 34, 2785–2792,doi: 10.1021/jasms.3c00298.

24. Xie, Y., De Luna Vitorino, F.N., Chen, Y., Lempiainen, J.K., Zhao, C., Steinbock, R.T., Lin, Z., Liu, X., Zahn, E., Garcia, A.L., et al. (2023) SWAMNA: a comprehensive platform for analysis of nucleic acid modifications. Chem Commun (Camb), 59, 12499–12502,doi: 10.1039/d3cc04402e.

25. Heiss, M., Borland, K., Yoluc, Y. and Kellner, S. (2021) Quantification of Modified Nucleosides in the Context of NAIL-MS. Methods Mol Biol, 2298, 279–306,doi: 10.1007/978-1-0716-1374-0_18.

26. Team, R.C. (2025). R Foundation for Statistical Computing, R, Vienna, Austria

27. Kerkhoff-Bernaciak, X. (2026), Zenodo, Vol. v1, pp. Code for Manuscript “Harmonized nucleoside mass spectrometry enables reproducible cross-platform RNA modification quantification” submission, 10.5281/zenodo.21028754.

28. Gu, H., Liu, G., Wang, J., Aubry, A.F. and Arnold, M.E. (2014) Selecting the correct weighting factors for linear and quadratic calibration curves with least-squares regression algorithm in bioanalytical LC-MS/MS assays and impacts of using incorrect weighting factors on curve stability, data quality, and assay performance. Anal Chem, 86, 8959–8966,doi: 10.1021/ac5018265.

29. Kellner, S., Neumann, J., Rosenkranz, D., Lebedeva, S., Ketting, R.F., Zischler, H., Schneider, D. and Helm, M. (2014) Profiling of RNA modifications by multiplexed stable isotope labelling. Chem Commun (Camb), 50, 3516–3518,doi: 10.1039/c3cc49114e.

30. Dal Magro, C., Keller, P., Kotter, A., Werner, S., Duarte, V., Marchand, V., Ignarski, M., Freiwald, A., Müller, R.-U., Dieterich, C., et al. (2018) A Vastly Increased Chemical Variety of RNA Modifications Containing a Thioacetal Structure. Angewandte Chemie International Edition, 57, 7893–7897,doi: 10.1002/anie.201713188.

31. (ICH), I.C.f.H. (2022)

32. Legrand, C., Tuorto, F., Hartmann, M., Liebers, R., Jacob, D., Helm, M. and Lyko, F. (2017) Statistically robust methylation calling for whole-transcriptome bisulfite sequencing reveals distinct methylation patterns for mouse RNAs. Genome Res, 27, 1589–1596,doi: 10.1101/gr.210666.116.

33. Wesseling, H. and Kaiser, S. (2025) Temporal and spatial profiling of ALKBH5 activity through NAIL-MS and compartmentalized RNA isolation. RNA, 31, 1589–1598,doi: 10.1261/rna.080593.125.

34. Richter, F., Plehn, J.E., Bessler, L., Hertler, J., Jorg, M., Cirzi, C., Tuorto, F., Friedland, K. and Helm, M. (2022) RNA marker modifications reveal the necessity for rigorous preparation protocols to avoid artifacts in epitranscriptomic analysis. Nucleic Acids Res, 50, 4201–4215,doi: 10.1093/nar/gkab1150.

35. Dong, M. and Dedon, P.C. (2006) Relatively small increases in the steady-state levels of nucleobase deamination products in DNA from human TK6 cells exposed to toxic levels of nitric oxide. Chem Res Toxicol, 19, 50–57,doi: 10.1021/tx050252j.

36. Zeng, W. and Bateman, K.P. (2023) Quantitative LC-MS/MS. 1. Impact of Points across a Peak on the Accuracy and Precision of Peak Area Measurements. J Am Soc Mass Spectrom, 34, 1136–1144,doi: 10.1021/jasms.3c00077.

37. Nelson, C.C. and McCloskey, J.A. (1992) Collision-induced dissociation of adenine. Journal of the American Chemical Society, 114, 3661–3668,doi: 10.1021/ja00036a014.

38. Jensen, S.S., Ariza, X., Nielsen, P., Vilarrasa, J. and Kirpekar, F. (2007) Collision-induced dissociation of cytidine and its derivatives. Journal of Mass Spectrometry, 42, 49–57,doi: 10.1002/jms.1136.

39. Gregson, J.M. and McCloskey, J.A. (1997) Collision-induced dissociation of protonated guanine. International Journal of Mass Spectrometry and Ion Processes, 165-**166**, 475–485,doi: 10.1016/S0168-1176(97)00163-8.

40. Nelson, C.C. and McCloskey, J.A. (1994) Collision-induced dissociation of uracil and its derivatives. Journal of the American Society for Mass Spectrometry, 5, 339–349,doi: 10.1016/1044-0305(94)85049-6.

41. Dumelin, C.E., Chen, Y., Leconte, A.M., Chen, Y.G. and Liu, D.R. (2012) Discovery and biological characterization of geranylated RNA in bacteria. Nature Chemical Biology, 8, 913–919,doi: 10.1038/nchembio.1070.

42. Kellner, S., Neumann, J., Rosenkranz, D., Lebedeva, S., Ketting, R.F., Zischler, H., Schneider, D. and Helm, M. (2014) Profiling of RNA modifications by multiplexed stable isotope labelling. Chemical Communications, 50, 3516–3518,doi: 10.1039/C3CC49114E.

43. Miyauchi, K., Kimura, S., Akiyama, N., Inoue, K., Ishiguro, K., Vu, T.-S., Srisuknimit, V., Koyama, K., Hayashi, G., Soma, A. et al. (2025) A tRNA modification with aminovaleramide facilitates AUA decoding in protein synthesis. Nature Chemical Biology, 21, 522–531,doi: 10.1038/s41589-024-01726-x.

44. Sun, J., Wu, J., Yuan, Y., Fan, L., Chua, Wei Lin P., Ling, Yan Han S., Balamkundu, S., Dwijapriya, Chay, Hazel Suen S., Begley, Thomas J., et al. (2025) tRNA modification profiling reveals epitranscriptome regulatory networks in Pseudomonas aeruginosa. Nucleic Acids Research, 53, gkaf696,doi: 10.1093/nar/gkaf696.

