## Supplemental Information for "Harmonized nucleoside mass spectrometry enables reproducible cross-platform RNA modification quantification"

^24^Brown University. Department of Molecular Biology, Cell Biology and Biochemistry, 70 Ship Street, Providence, RI 02903, United States

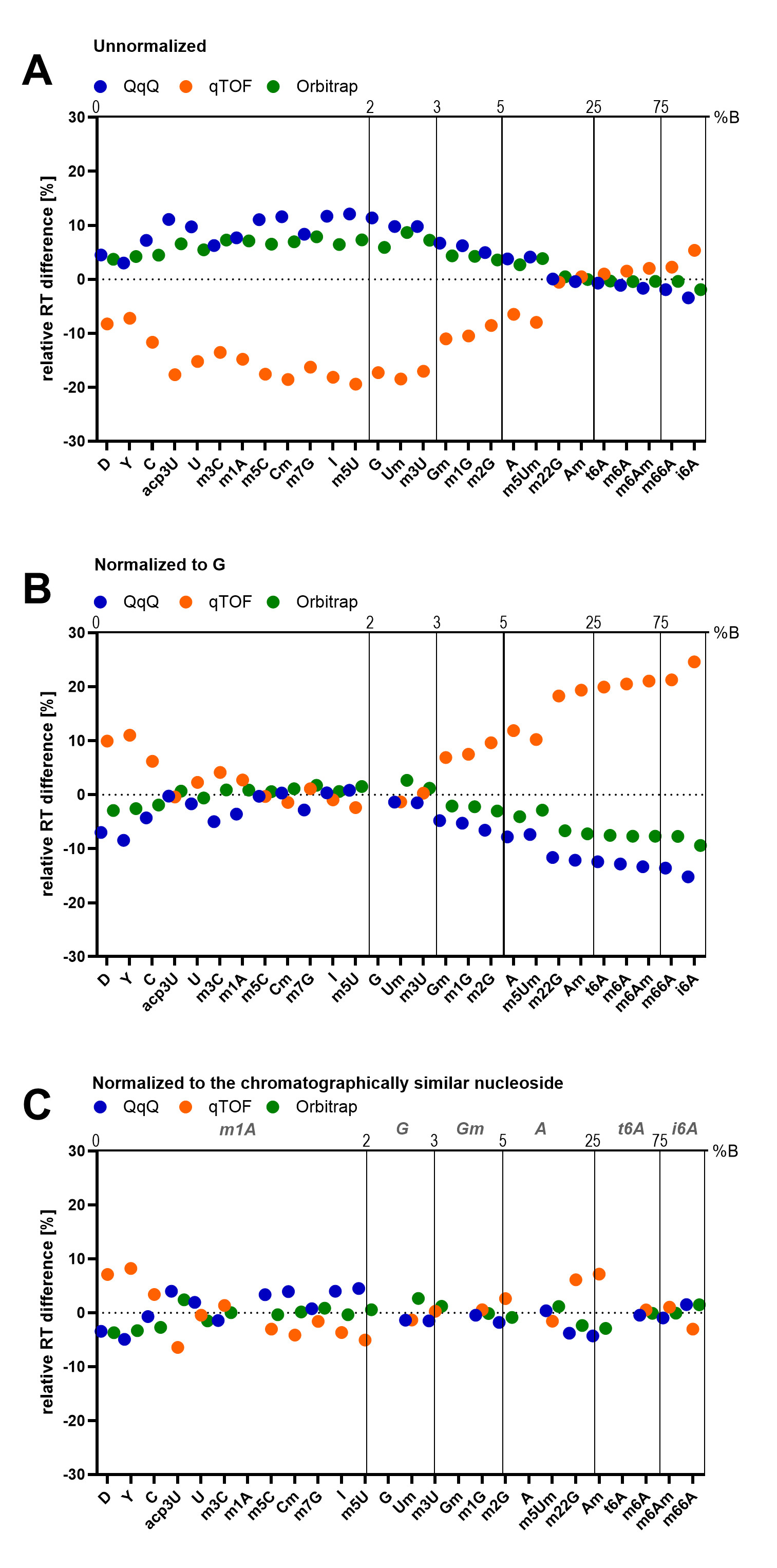

**Figure S1** Relative retention time differences calculated from the mean retention time of all systems. The upper y-axis indicates the gradient solvent composition, with the grid indicating changes in solvent composition at the mean retention time of the corresponding modification. **(A)** Relative differences of unnormalized retention times. **(B)** Relative differences of retention times normalized to G. **(C)** Relative differences of retention times normalized to the chromatographically similar nucleoside. The text in cursive indicates which nucleosides were used as a reference for the respective chromatographic section.

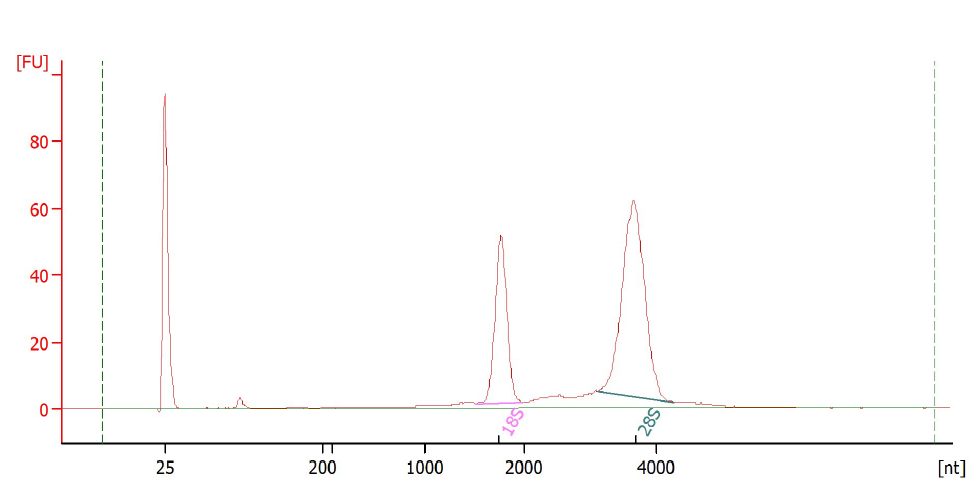

**Figure S2** Electropherogram of small-RNA depleted RNA isolated from Human B-cells (GM12878) analyzed using the Agilent 2100 Bioanalyzer and the RNA 6000 Pico Kit. The prominent peaks correspond to the 18S and 28S ribosomal RNA subunits.
